# Modeling the effects of vaccination against multiple strains of porcine reproductive and respiratory syndrome at the barn level

**DOI:** 10.1101/2024.08.06.606602

**Authors:** Faith Kennedy, Nicolas Cardenas, Jason A. Galvis, Cesar Corzo, Gustavo Machado

## Abstract

Porcine reproductive and respiratory syndrome (PRRSV) remains costly for the swine industry. Measures intended to irradicate or reduce the prevalence of PRRSV have limited effectiveness. Vaccination, the primary control strategy for PRRSV, has yet to be thoroughly investigated to ensure an optimal strategy is being implemented. Due to insufficient cross-immunity between strains, it is prudent to administer the most influential vaccine for the most dominant or threatening PRRSV strain in circulation at a given time. With this in mind, we demonstrated the benefits of targeted vaccination using metapopulation-based modeling. By using up-to-date data describing real farms to simulate the effects of variable vaccine efficacy in both a single and a double strain system, we showed that tailoring vaccines to prevalent PRRSV strains reduces infections, but using multiple vaccines may be more effective if strains vary in vaccine response. Therefore, the impact of the strain-specific effectiveness of market vaccines, paired with computational methods that allow tracking the effects on PRRVS dissemination at individual barns.

## Introduction

Porcine reproductive and respiratory syndrome virus (PRRSV) is a significant player in the disease landscape of commercial swine production in North America (Valdes-Donoso & Jarvis, 2022) and other major swine-producing countries (Zhang et al., 2022). Losses from the illness in the U.S. are believed to surpass $660 million annually (Joltkamp et al., 2013; Neumann et al., 2005). PRRSV poses significant risks to sow farms, mainly by causing reproductive failure (Hopper et al., 1992; Karniychuk et al., 2012; Terpstra et al., 1991). Additionally, illness in developing pigs can impede growth, increase time to slaughter, and raise the risk of death (Schweer et al., 2016). Mitigating the spread of PRRSV has the potential to alleviate the financial strain on the pork industry greatly. *In silico* methods allow for an in-depth investigation of the introduction and dissemination of PRRSV and the prediction of best practices for control strategies (Barh et al., 2020).

Several PRRSV strains traverse inter-farm networks within the U.S. (Kikuti et al., 2021; Shi et al., 2010). Some evidence suggests that the dominant strain of PRRSV changes approximately every three years (Paploski et al., 2021). While vaccination against PRRSV is commonly utilized in the swine industry, little is known about the impact of targeted vaccination on the ability of individual PRRSV strains to permeate swine populations (Bitsouni et al., 2019). There is mixed evidence in the literature suggesting there may not be substantial cross-immunity between strains (Madapong et al., 2020; Chae, 2021); this is especially troubling given that current market vaccines are insufficient to properly control PRRSV (Hu & Zhang, 2014; Renukaradhya et al., 2015; Vu et al., 2017). Being able to match vaccine choices to the specific strains that are at the highest risk has the potential to boost immunity in a much-needed way. Therefore, strain-specific vaccination may be necessary to reduce PRRSV circulation significantly.

PRRSV strain classification follows restriction fragment length polymorphism (RFLP) or lineage designations. However, RFLP appears insufficient to distinguish different variants for predictive analysis and vaccine strategization (Sang-Ho et al., 2004; Shi et al., 2010; Trevisan et al., 2021). RFLPs are based solely on the ORF5 site of PRRSV, which is not considered an active site in the virus and is associated with high levels of mutation (Darwich et al., 2011; Mardassi et al., 1995; Sang-Ho et al., 2004). Therefore, classification based on RFLP is limited in predicting functional changes in the virus (Sang-Ho et al., 2004; Shi et al., 2010; Trevisan et al., 2021).

Lineage classifications take the evolutionary history of the virus into account and are more likely to have real biological meaning, making it the preferred method for strain classification. Using lineages for PRRSV classification is especially important, considering that the level of cross-immunity between strains may be unsubstantial. Determining evolutionary relationships and narrowing down phylogenetic groupings more precisely could help predict the spread of individual variants and better estimate the effects of vaccination.

Over the years, several compartmental models have been developed to describe the spread of PRRSV, used to investigate the spread of PRRSV between farms (Arruda et al., 2017; Galvis, Corzo, & Machado, 2022; Galvis, Corzo, Prada, et al., 2022; Thakur et al., 2015; Valdes-Donoso & Jarvis, 2022), while others focus on metapopulation dynamics (Arruda et al., 2016; Bitsouni et al., 2019; Evans et al., 2010; Jeong et al., 2014; Phoo-ngurn et al., 2019; Zou et al., 2020). A susceptible-infected-recovered (SIR) or susceptible-undetected-detected (SUD) model was commonly used. For many of the models that consider within-farm dynamics, the studied populations were kept closed or included little movement in and out of the herd (Bitsouni et al., 2019; Evans et al., 2010; Phoo-ngurn et al., 2019; Zou et al., 2020). This dynamic is unlikely to accurately represent the population dynamics of commercial swine farms in North America, which are vertically integrated and high levels of animals of different sources mixing at wean-to-finisher farms (Boripun et al., 2021; Cardenas et al., 2024). While some within-farm PRRSV transmission models implemented control actions such as herd closure (Arruda et al., 2016; Jeong et al., 2014), gilt acclimatization (Jeong et al., 2014), and vaccination (Arruda et al., 2016; Jeong et al., 2014), only a 2019 paper by Bitsouni et al modeled the effects of variable vaccine effectiveness. However, their simulations feature one vaccine at a time used to combat a single unified strain of the virus and do not consider the direct impact of genetic variation on vaccine efficacy.

To address the role of genetic diversity in continued PRRSV outbreaks, we sought to investigate the benefit of strain-specific vaccination. Using real-world data, we determined the most common PRRSV lineages circulating in recent years. We incorporated actual swine movements into our model, promoting the accuracy and applicability of our results. In addition to this information, our new model incorporates multiple strains and multiple vaccines with varying levels of efficacy.

## Methods

### Population data

This study used data from 98 commercial swine farms in one U.S. state. Our database includes the number of barns per farm extracted from the Rapid Access Biosecurity application (RABapp™) database (Machado et al., 2023). Briefly, The RABapp™ results from a multi-state consortium of academic researchers, animal health officials, and swine producers that serves as a platform for standardizing the approval of Secure Pork Supply (SPS) biosecurity plans and as a centralized database for animal, semen, and vehicle movement data. An SPS biosecurity plan comprises a written section and a farm map of fourteen spatial features, including the line of separation (LOS)(Supplementary Material Figure S1). Each premises map has at least one LOS established as a control boundary to prevent pathogen movement into pig housing areas (a.k.a. barns). Here, we used LOS counts to generate a database with the number of barns per farm. In addition, the RABapp™ database also includes the total number of pigs per farm, here; here, we divided the total number of pigs by the number of barns per farm to generate barn-level initial pig populations. With the starting population of each barn, we used the movement of pigs to update the farm’s population; for instance, farm *i* with a pig capacity of 1,200 pigs, a day *t* farm *i* received one load of 400 pigs, and on day *t+1* received three loads of 400 pigs. On day t + 180 days two loads of 400 pigs were shipped to a slaughterhouse, and finally, on day t+190, the remaining pigs were shipped, and the site was empty until the next load of pigs was received on day t + 193.

### PRRSV sequence data

Our data set comprises 340 ORF-5 sequences collected from July 2022 until December 2023, obtained from three commercially unrelated pig production systems located in North Carolina, United States. Sequences were generated through outbreak investigation and from regular surveillance activities performed by each production system and shared with the Morrison Swine Health Monitoring Program (MSHMP) (Perez et al., 2019). PRRSV sequences were constructed into a phylogenetic tree using a k-nearest-neighbors algorithm, and unknown lineages were assigned. The resulting phylogeny was confirmed to be accurate with R^2^ = 0.98 and can be seen in Supplementary Material Figure S2. Of the four lineages found in the data since 2020, lineage L1A was found to be the most prevalent, with a 75% majority.

### Model structure

To model the spread of PRRSV on wean-to-finish farms at the barn level, we defined the following system of differential equations according to the compartmental dynamics shown in Figure 1, where the stated parameters are as outlined in Table 1

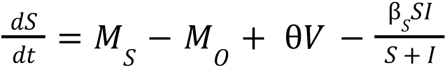

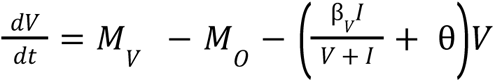

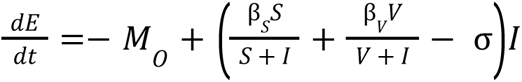

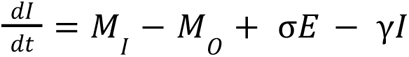

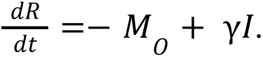

**Figure 1.**
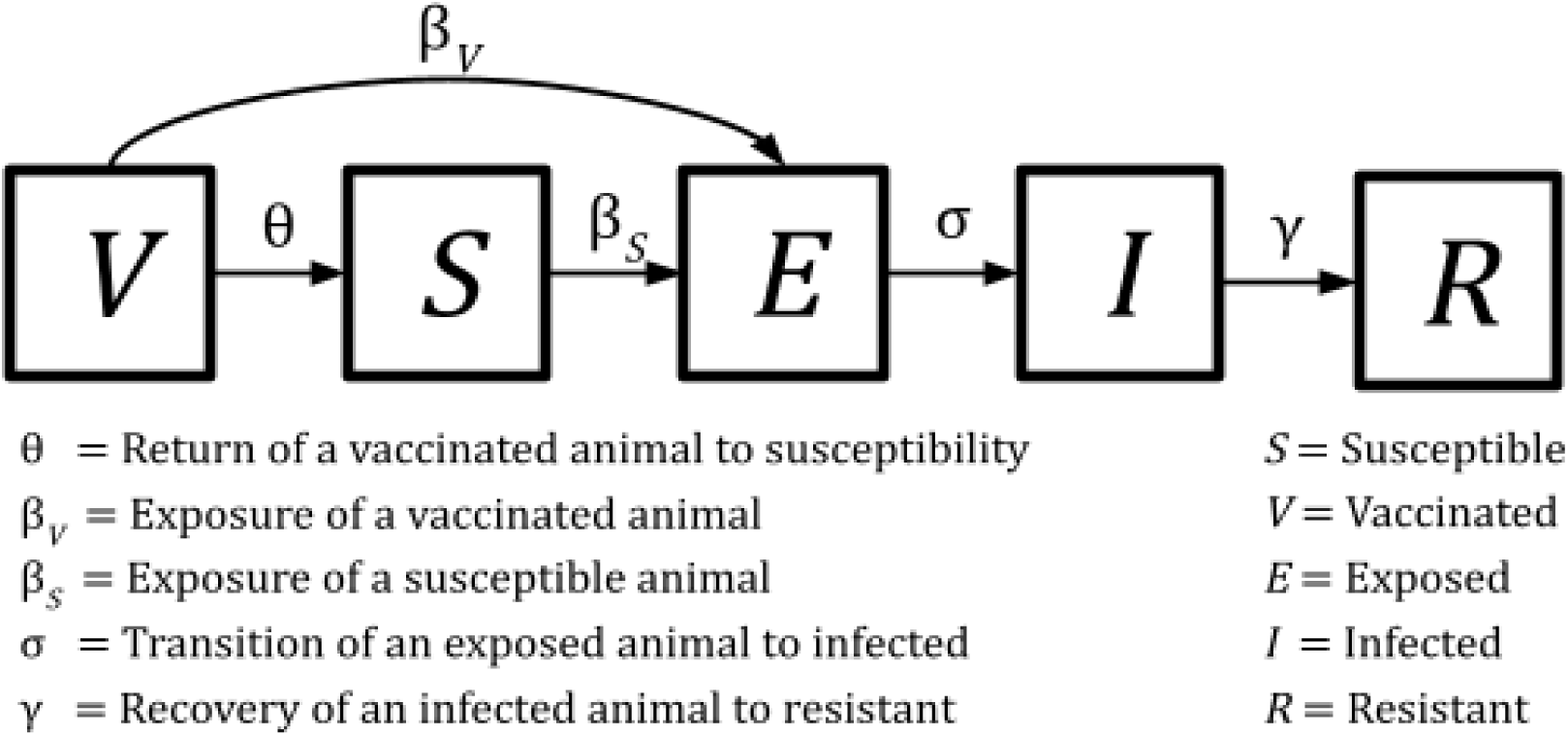
Compartmental System: The diagram illustrates the system used in model simulations. Each boxed letter corresponds to the compartment indicated by the legend. Movements between compartments are according to the rates given in Table 1.

**Table 1.** Summary of parameters used in the disease dynamics model.

| Parameter | Rate at individuals per day | Description | Reference |
| --- | --- | --- | --- |
| $\beta_s$ | 0.2 | Infection of a Susceptible | (Pileri et al., 2015) |
| $\beta_v$ | 0.07, 0.13 | Infection of a Vaccinated | Calculated using efficacy level and $\beta_s$ |
| $\sigma$ | 1/1.5 | Transition to Infection | (Pileri et al., 2015) |
| $\gamma$ | 1/112 | Recovery of Infected | (Angulo et al., 2023) |
| $\theta$ | 1/133 | Return of Vaccinated to Susceptible | (Zoetis Services, 2021) |
| $M_s$ | From data | Movement to Susceptible | - |
| $M_o$ | From data | Movement out | - |
| $M_v$ | From data | Movement to Vaccinated | - |
| $M_i$ | From data | Movement to Infected | - |

Our model includes vaccination at the barn level, which is assumed to provide incomplete immunity where vaccinated individuals are exposed at a rate β_*V*_ upon contact with an infected individual with barns. Vaccination is considered to wain over time, and individuals return to susceptibility at a rate θ. Susceptible individuals are assumed to become exposed at a rate of β_s_ upon contact with an infected individual. Exposed individuals become infectious at a rate σ. Infected individuals develop resistance at a rate γ. We estimate contact rates at the barn level using the product of the two populations scaled by the total size of the populations involved. We assumed that individual pigs were always homogeneously mixed and evenly distributed within the barns. Inward movements are denoted *M*_*S*_, *M*_V_, and *M*_I_ according to the corresponding compartment, and outward movements are denoted *M_O_* (Table 1). Outward movements for each compartment were scaled according to the proportion of the population represented.

In total, simulations were performed for 98 farms, all part of one production system in North Carolina. In lieu of market movement, every farm population was assumed to follow an “all-in, all-out”. A gap between incoming shipments of over 30 days was considered to indicate a “turnover event” where all remaining pigs were removed, and the barn(s) were cleaned, thus all remaining individuals were removed the day prior to the new movement. Results were simulated over the time period from July 1, 2022 through December 31, 2023 and aligned according to the first turnover event during this time period (Supplementary Material Figure S3). For our simulation scenarios, each shipment of pigs was assumed to arrive 90% vaccinated or 90% susceptible as appropriate for the scenario, with the remaining 10% infected pigs (Supplementary Material Figure S4). In the case of multiple vaccine scenarios, the 10% infected population was split into 7.5% strain 1 and 2.5% strain 2 (Supplementary Material Figure 4B), in accordance with our phylogenetic data (Supplementary Material Figure S2).

We simulated eight scenarios that featured the dynamics of one and two strains and vaccination scenarios with one and two strain-specific vaccines. For the one-strain scenarios, vaccine one was assumed to be 33% effective and vaccine two was 67% effective. In the two strain scenarios, efficacy was assumed to vary for different vaccine-strain pairings, thus, efficacy was defined as outlined in Table 2b. Vaccine one was considered to be 67% effective for Strain one and 33% effective for Strain two, while Vaccine two was considered to be 33% effective for Strain one and 67% effective for Strain two. The infection rates were calculated using the expression 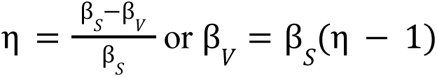, where η is the vaccine’s efficacy.

**Table 2:** Description of the different scenarios simulated. (top) gives information about the strains (S1 and S2) and vaccines (V1 and V2) used, while (bottom) provides the efficacy levels used.

| Scenario | # Strains | # Vaccines | S1:S2 Ratio | V1:V2 Ratio |
| --- | --- | --- | --- | --- |
| 1 | 1 | No vaccine | 1:0 | No vaccine |
| 2 | 1 | 1 | 1:0 | 1:0 |
| 3 | 1 | 1 | 1:0 | 0:1 |
| 4 | 1 | 2 | 1:0 | 1:1 |
| 5 | 2 | No vaccine | 3:1 | No vaccine |
| 6 | 2 | 1 | 3:1 | 1:0 |
| 7 | 2 | 2 | 3:1 | 1:1 |
| 8 | 2 | 1 | 3:1 | 0:1 |

| | Efficacy ($\eta$) | |
| --- | --- | --- |
|  | Vaccine 1 (V1) | Vaccine 2 (V2) |
| Strain 1 (S1) | 67% | 33% |
| Strain 2 (S2) | 33% | 67% |

## Results

Observing the dynamics of our model on a fixed population, we see that the model achieves a dynamic consistent with the existing data (Supplementary Material Figure S5). However, the number of cases arising from the two strain scenarios is underestimated when the fixed population is compared to the real data (Figure 2; Supplementary Material Figure S5). Incorporating ingoing and outgoing movements also shows that PRRSV outbreaks are unlikely to be cleared before each cohort is moved to the market (Figure 2).

**Figure 2.**
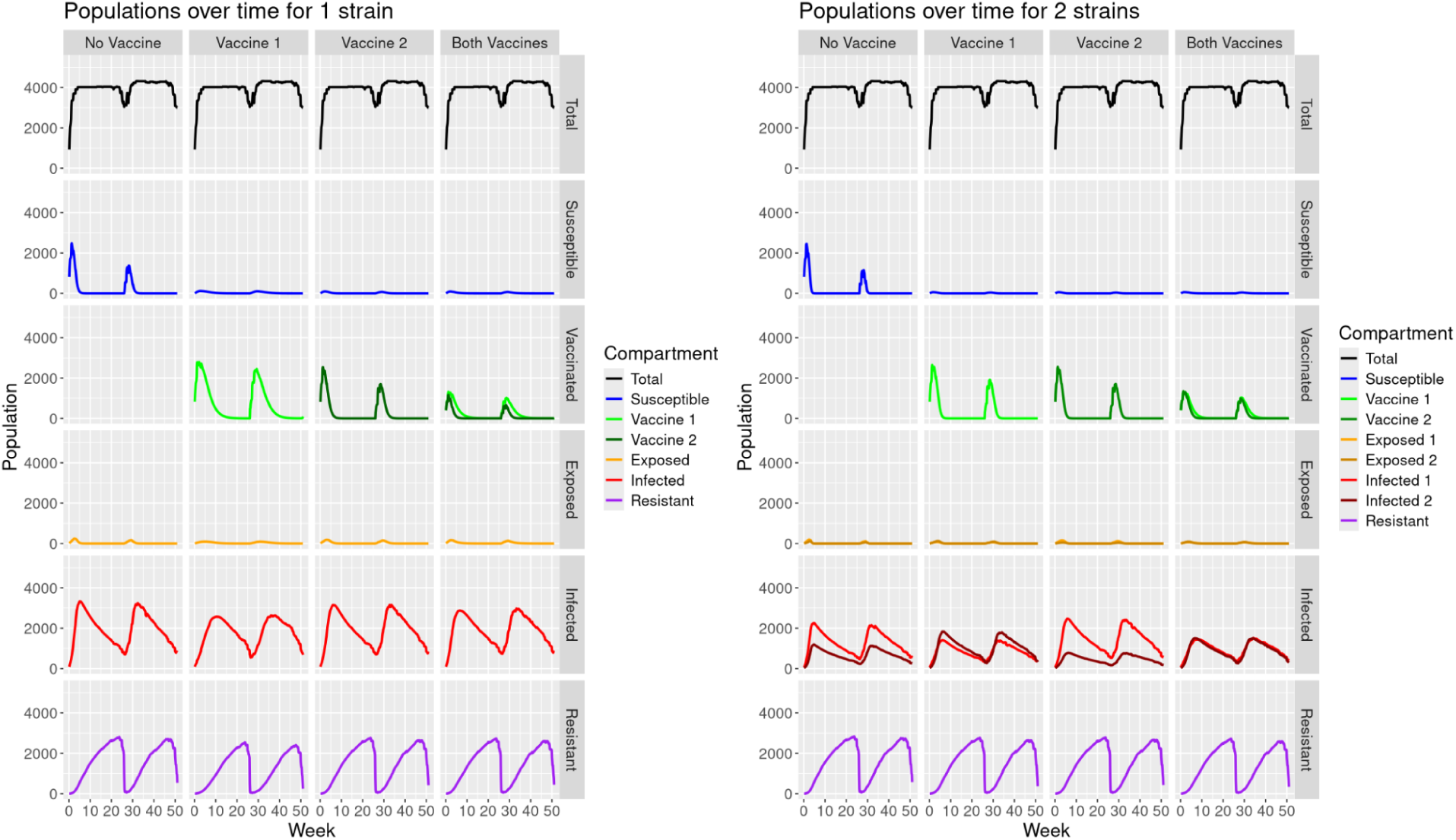
shows the Effect of the PRRV vaccine’s uncertainty on barn-level dissemination. The left panel shows the dynamics of the circulation of one PRRV strain, and the right panel shows the dissemination of two strains and the effects of vaccination.

The PRRSV prevalence for the model with one strain peaks at a much lower value when a more effective vaccine is used. This is especially true when the more effective vaccine is used exclusively, but it is still true to some extent when a mixture of vaccines is used. However, these dynamics change when multiple strains are present. In the model with two strains, vaccinating for only one strain in the global population ultimately resulted in similar numbers of relative infections regardless of the vaccine used (Figure 3). It is clear that using both vaccines resulted in the lowest number of related infections. In all cases, vaccination lowered the peak proportion of infections compared to a baseline of no immunization (Figure 3).

**Figure 3:**
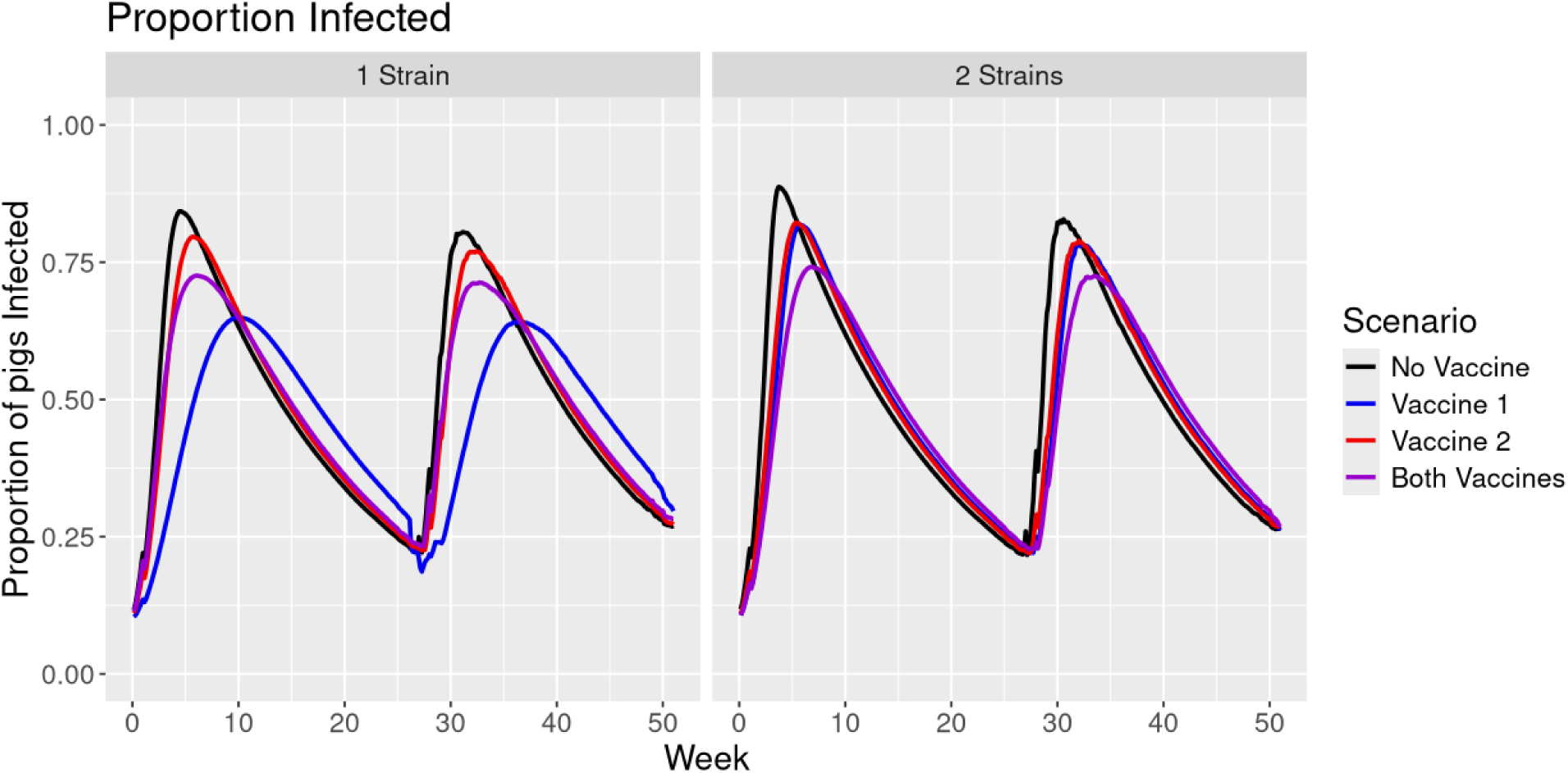
The median proportion of pigs infected over time across all farms aligned by the first incoming shipment of 2023.

## Discussion

PRRSV near global distribution has caused significant losses to the pork industry, and today, its control is mainly driven by the use of pharmacological control actions, which can be preventative or reactive vaccination (Galvis, Corzo, Prada, et al., 2022). While the effectiveness of vaccination over epidemics has yet to be explored in detail using a mechanistic model at the farm level (Galvis, Corzo, Prada, et al., 2022), barn-level examinations still need to be explored. The role of the circulation of multiple PRRSV strains and the population-level effect of vaccinating to specific PRRSV lineage needs to be better understood. To that end, we have reconstructed the pig population dynamics at the barn level to model the dissemination of single and multi-PRRSV staging dissemination. Our results demonstrated that the model confirmed the delay of PRRSV peaks when the vaccine matched the single strain circulation and the lack of significant effects over the dissemination when more than one strain is present.

While population dynamics simulation is an essential part of epidemiology, there are limits to the exploration that can be done using solely simulated data. Swine production generally follows consistent and predictable patterns that can be incorporated into population dynamics models to increase accuracy and drive new findings. Increased access to data describing movement, environment, and disease within and between farms is valuable in formulating more useful and applicable models going forward. There is mounting evidence to suggest that genetic variation within the PRRSV species impacts how the virus spreads. Our results suggest that this dynamic should be carefully considered when determining best practices and control strategies at the barn level. It is difficult to predict the true benefit of targeted vaccination without first increasing our knowledge of how different strains are impacted by specific vaccines. It is not yet understood how or why different strains may be more virulent or whether specific strains are responsible for more severe symptoms. It is also unclear whether certain vaccines provide greater cross-protection than others or whether the administration of multiple vaccines in the same individual can provide stacked immunity to multiple strains. As this information is uncovered, it will become easier to alter existing models to predict which vaccines should be administered in real time. Future works may combine the disease transmission model and phylogenetic analysis to increase resolution for identifying dominant strains in the field. Integrating the existing model into an inter-farm network to model the spread of the virus between farms would be crucial in determining how strains are transmitted on a larger scale.

## Conclusion

We developed a mathematical model to investigate the spread of multiple PRRSV strains within commercial swine barns, incorporating multiple vaccines with varying degrees of effectiveness. First, we estimated the most common lineages and their relative prevalence for use in our simulations. An assessment of the model dynamics supports the validity of model parameters in addition to some identifiable differences when real movement data is used. We showed that tailoring vaccine administration to the most prevalent strains results in reduced infection when PRRSV is considered a single unified virus, but introducing multiple vaccines into the population may be more beneficial if vaccine efficacy varies among different strains. These results show that understanding how specific strain-vaccine pairings interact and using this to inform real-life vaccination decisions would be a powerful tool in reducing the incidence of PRRSV.

## Conflict of interest

All authors confirm that there are no conflicts of interest to declare.

## Ethical statement

The authors confirm the journal’s ethical policies, as noted on the journal’s author guidelines page. Ethics permits were unnecessary since this work did not involve animal sampling or questionnaire data collection by the researchers.

## Data Availability Statement

The data supporting this study’s findings are not publicly available and are protected by confidential agreements; therefore, they are not available.

## Funding

This work was supported by Inter-Disciplinary Engagement in Animal Systems (IDEAS), 2022-68014-37266 from the USDA National Institute of Food and Agriculture.

## Acknowledgements

The authors would like to thank the production system that worked with us for this study.

## Notes

### Competing Interest Statement

The authors have declared no competing interest.

### Summary of Updates

adding funding to the list of acknolegments

## Bibliography

1. Angulo, J., Yang, M., Robira, A., Davies, P. R., & Montserrat, T. (2023). Infection dynamics and incidence of wild-type porcine reproductive and respiratory syndrome virus in growing pig herds in the U.S. Midwest. Preventive Veterinary Medicine, 217.

2. Arruda, A. G., Friendship, R., Carpenter, J., Greer, A., & Poljak, Z. (2016). Evaluation of Control Strategies for Porcine Reproductive and Respiratory Syndrome (PRRS) in Swine Breeding Herds Using a Discrete Even Agent-Based Model. PLoS ONE, 11(11). 10.1371/journal.pone.0166596

3. Arruda, A. G., Poljak, Z., Knowles, D., & McLean, A. (2017). Development of a stochastic agent-based model to evaluate surveillance strategies for detection of emergent porcine reproductive and respiratory syndrome. BMC Veterinary Research, 13(171).

4. Barh, D., Yiannakopoulou, E. C., Salawu, E. O., Battacharjee, A., Chowbina, S., Nalluri, J. J., Ghosh, P., & Azevedo, V. (2020). In silico disease model: From simple networks to complex diseases. Animal Biotechnology, 2020, 441–460.

5. Bitsouni, V., Lycett, S., Opriessnig, T., & Doeschl-Wilson, A. (2019). Predicting vaccine effectiveness in livestock populations: A theoretical framework applied to PRRS virus infections in pigs. PLoS ONE, 14(8). e0220738

6. Boripun, R., Mitsuwan, W., Kulnanan, P., Thomrongsuwannakij, T., & Kitpipit, W. (2021). Analysis of culling reasons during the breeding cycle and lifetime performance: The strategy to remove crossbred Landrace and Large White sows under tropical climate. Veterinary World, 14(12), 3170–3174.

7. Cardenas, N. C., Valencio, A., Sanchez, F., O’Hara, K. C., & Machado, G. (2024). Analyzing the intrastate and interstate swine movement network in the United States. Preventive Veterinary Medicine, 106264. 10.1016/j.prevetmed.2024.106264

8. Chae, C. (2021). Commercial PRRS Modified-Live Virus Vaccines. Vaccines (Basel), 9(2), 185. 10.3390/vaccines9020185

9. Darwich, L., Gimeno, M., Sibila, M., Diaz, I., Torre, E. de la, Dotti, S., Kuzemtseva, L., Martin, M., Pujols, J., & Mateu, E. (2011). Genetic and immunobiological diversities of porcine reproductive and respiratory syndrome genotype I strains. Veterinary Microbiology, 150(1–2), 49–62.

10. Evans, C. M., Medley, S. J., Creasey, S. J., & Green, L. E. (2010). A stochastic mathematical model of the within-herd transmission dynamics of porcine reporductive and respiratory syndrome virus (PRRSV): Fade-out and persistence. Preventive Veterinary Medicine, 93(4), 248–257.

11. Galvis, J. A., Corzo, C. A., & Machado, G. (2022). Modelling and assessing additional transmission routes for porcine reproductive and respiratory syndrome virus: Vehicle movements and feed ingredients. Transboundary and Emerging Diseases, 69(5), 2405–3143.

12. Galvis, J. A., Corzo, C. A., Prada, J. M., & Machado, G. (2022). Modelling the transmission and vaccination strategy for porcine reproductive and respiratory syndrome virus. Transboundary and Emerging Diseases, 69, 485–500.

13. Hopper, S. A., White, M. E., & Twiddy, N. (1992). An outbreak of blue-eared pig disease (porcine reproductive and respiratory syndrome) in four pig herds in Great Britain. The Veterinary Record, 131(7), 140–144. 10.1136/vr.131.7.140

14. Hu, J., & Zhang, C. (2014). Porcine Reproductive and Respiratory Syndrome Virus Vaccines: Current Status and Strategies to a Universal Vaccine. Transboundary and Emerging Diseases, 61, 109–120.

15. Jeong, J., Aly, S. S., Cano, J. P., Polson, D., Kass, P. H., & Perez, A. M. (2014). Stochastic model of porcine reproductive and respiratory syndrome virus control strategies on a swine farm in the United States. American Journal of Veterinary Research, 75(3). 10.2460/ajvr.75.3.260

16. Joltkamp, D. J., Kliebenstein, J. B., Neumann, E. J., Zimmerman, J. J., Rotto, H. F., Yoder, T. K., Wang, C., Yeske, P. E., Mowrer, C. L., & Haley, C. A. (2013). Assessment of the economic impact of porcine reproductive and respiratory syndrome virus on United States pork producers. Journal of Swine Health and Production, 21(2), 72–84. 10.31274/ans_air-180814-28

17. Karniychuk, U. U., Saha, D., Vanhee, M., Geldhof, M., Cornillie, P., Caij, A. B., Regge, N. D., & Nauwynck, H. J. (2012). Impact of a novel inactivated PRRS virus vaccine on virus replication and virus-induced pathology in fetal implantation sites and fetuses upon challenge. Theriogenology, 78(7), 1527–1537.

18. Kikuti, M., Sanhueza, J., Vilalta, C., Paploski, I. A. D., VanderWaal, K., & Corzo, C. A. (2021). Porcine reproductive and respiratory syndrome virus 2 (PRRSV-2) genetic diversity and occurrence of wild type and vaccine-like strains in the United States swine industry. PLoS ONE, 16(11). 10.1371/journal.pone.0259531

19. Machado, G., Galvis, J., Freeman, A., Sanchez, F., Fleming, C., Hong, X., Mills, K., & Sykes, A. (2023). The Rapid Access Biosecurity (RAB) app^TM^ Handbook. 10.17605/OSF.IO/Z5WBJ

20. Madapong, A., Saeng-chuto, K., Boonsoongnern, A., Tantituvanont, A., & Nilubol, D. (2020). Cell-mediated immune response and protective efficacy of porcine reproductive and respiratory syndrome virus modified-live vaccines against co-challenge with PRRSV-1 and PRRSV-2. Scientific Reports, 10. 10.1038/s41598-020-58626-y

21. Mardassi, H., Mounir, S., & Dea, S. (1995). Structural Gene Analysis of a Quebec Reference Strain of Porcine Reproductive and Respiratory Syndrome Virus (PRRSV). In Corona– and Related Viruses (Vol. 380, pp. 277–281). https://pubmed.ncbi.nlm.nih.gov/8830492/

22. Neumann, E. J., Kliebenstein, J. B., Johnson, C. D., Mabry, J. W., Bush, E. J., Seitzinger, A. H., Green, A. L., & Zimmerman, J. J. (2005). Assessment of the economic impact of porcine reproductive and respiratory syndrome on swine production in the United States. Journal of the American Veterinary Medicine Association, 227(3), 385–392. 10.2460/javma.2005.227.385

23. Paploski, I. A. D., Pamornchainavakul, N., Makau, D. N., Rovira, A., Corzo, C. A., Schroeder, D. C., Cheeran, M. C.-J., Doeschl-Wilson, A., Kao, R. R., Lycett, S., & VanderWaal, K. (2021). Phylogenetic Structure and Sequential Dominance of Sub-Lineages of PRRSV Type-2 Lineage 1 in the United States. Vaccines, 9(6), 608.

24. Perez, A. M., Linhares, D. C. L., Arruda, A. G., VanderWaal, K., Machado, G., Vilalta, C., Sanhueza, J. M., Torrison, J., Torremorell, M., & Corzo, C. A. (2019). Individual or Common Good? Voluntary Data Sharing to Inform Disease Surveillance Systems in Food Animals. Frontiers in Veterinary Science, 6, 194. 10.3389/fvets.2019.00194

25. Phoo-ngurn, P., Kiataramkul, C., & Chamchod, F. (2019). Modeling the spread of porcine reproductive and respiratory syndrome virus (PRRSV) in a swine population: Transmission dynamics, immunity information, and optimal control strategies. Advances in Continuous and Discrete Models, 2019(432). 10.3389%2Ffvets.2021.727895

26. Pileri, E., Gilbert, E., Soldevila, F., Garcia-Saenz, A., Pujols, J., Diaz, I., Darwich, L., Casal, J., Martin, M., & Mateu, E. (2015). Vaccination with a genotype 1 modified live vaccine against porcine reproductive and respiratory syndrome virus significantly reduces viremia, viral shedding and transmission of the virus in a quasi-natural experimental model. Veterinary Microbiology, 175(1), 7–16. 10.1016/j.vetmic.2014.11.007

27. Renukaradhya, G. J., Meng, X. J., Calvert, J. G., Roof, M., & Lager, K. M. (2015). Live porcine reproductive and respiratory syndrome virus vaccines: Current status and future direction. Vaccine, 33(33), 4069–4080.

28. Sang-Ho Cha, Chih-Cheng Chang, & Kyoung-Jin Yoon. (2004). Instability of the restriction fragment length polymorphism pattern of open reading frame 5 of porcine reproductive and respiratory syndrome virus during sequential pig-to-pig passages. Journal of Clinical Microbiology, 42(10), 4462–4467. 10.1128/JCM.42.10.4462-4467.2004

29. Schweer, W. P., Schwartz, K., Burrough, E. R., Yoon, K. J., Sparks, J. C., & Gabler, N. K. (2016). The effect of porcine reproductive and respiratory syndrome virus and porcine epidemic diarrhea virus challenge on growing pigs I: Growth performance and digestibility. Journal of Animal Science, 94(2), 514–522. 10.2527/jas.2015-9834

30. Shi, M., Lam, T. T.-Y., Hon, H.-C., Hui, R. K.-H., Faaberg, K. S., Wennblom, T., Murtaugh, M. P., Stadejek, T., & Leung, F. C.-C. (2010). Molecular epidemiology of PRRSV: A phylogenetic perspective. Virus Research, 154(1–2), 7–17. 10.1016/j.virusres.2010.08.014

31. Terpstra, C., Wensvoort, G., & Pol, J. M. (1991). Experimental reproduction of porcine epidemic abortion and respiratory syndrome (mystery swine disease) by infection with Lelystad virus: Koch’s postulates fulfilled. The Veterinary Quarterly, 13(3), 131–136. 10.1080/01652176.1991.9694297

32. Thakur, K. K., Sanchez, J., Hurnik, D., Poljak, Z., Opps, S., & Revie, C. W. (2015). Development of a network based model to simulate the between-farm transmission of the porcine reproductive and respiratory syndrome virus. Veterinary Microbiology, 180(3–4), 212–222.

33. Trevisan, G., Sharma, A., Gauger, P., Harmon, K. M., Zhang, J., Main, R., Zeller, M., Linhares, L. C. M., & Linhares, D. C. L. (2021). PRRSV2 genetic diversity defined by RFLP patterns in the United States from 2007 to 2019. Journal of Veterinary Diagnostic Investigation, 33(5), 920–931. 10.1177/10406387211027221

34. Valdes-Donoso, P., & Jarvis, L. S. (2022). Combining epidemiology and economics to assess control of a viral endemic animal disease: Porcine Reproductive and Respiratory Syndrome (PRRS). PLoS ONE, 17(9).

35. Vu, H. L. X., Pattnaik, A. K., & Osorio, F. A. (2017). Strategies to broaden the cross-protective efficacy of vaccines against porcine reproductive and respiratory syndrome virus. Veterinary Microbiology, 206, 29–34.

36. Zhang, Z., Li, Z., Li, H., Yang, S., Ren, F., Bian, T., Sun, L., Zhou, B., Zhou, L., & Qu, X. (2022). The economic impact of porcine reproductive and respiratory syndrome outbreak in four Chinese farms: Based on cost and revenue analysis. Frontiers in Veterinary Science, 9. 10.3389/fvets.2022.1024720

37. Zoetis Services. (2021). Fostera PRRS: Porcine reproductive & respiratory syndrome vaccine.

38. Zou, J., Upadhyay, R. K., Pratap, A., & Zhang, Z. (2020). Dynamics of a delayed SIR model for the transmission of PRRSV among a swine population. Advances in Continuous and Discrete Models, 2020(351). 10.1186/s13662-020-02814-7

