## Supplementary material for "Modeling the effects of vaccination against multiple strains of porcine reproductive and respiratory syndrome at the barn level": ss

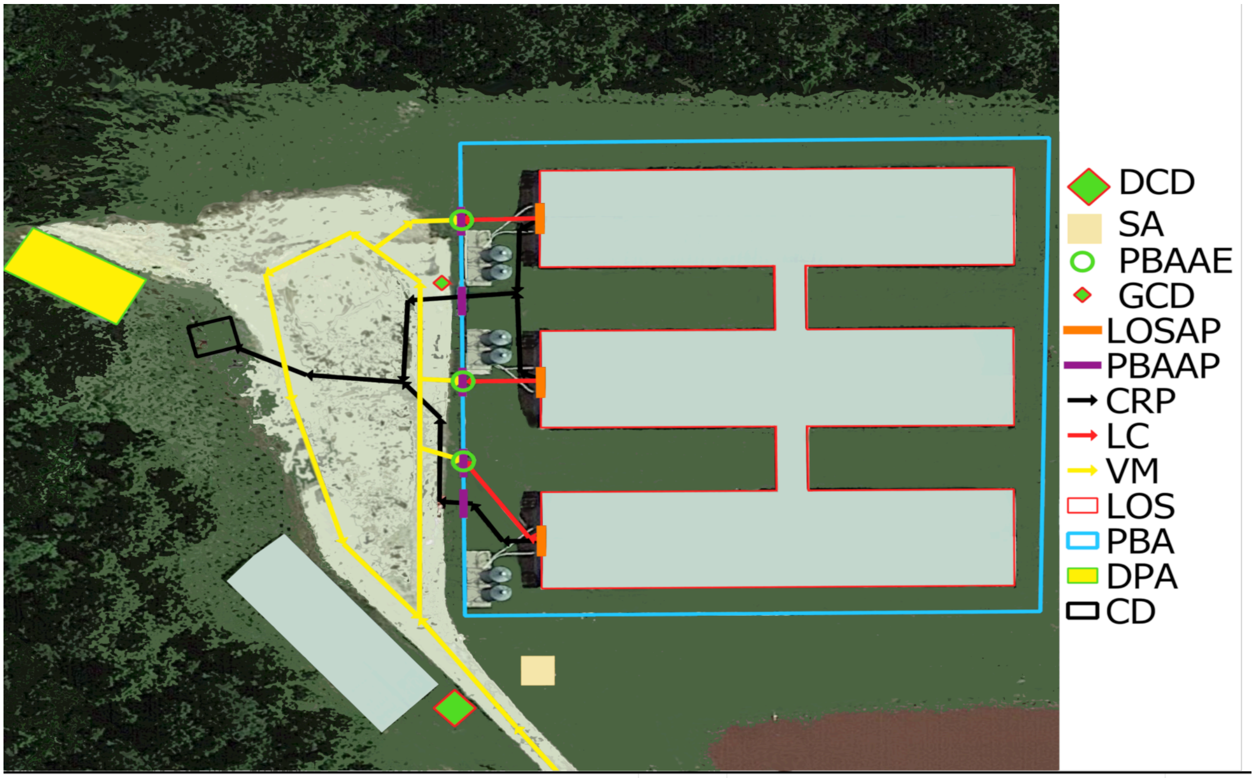


| Legend | Symbol | Legend | Symbol |
|---|---|---|---|
| LOSAE (LOS Animal Emergency) | | PBAAP (PBA Access Point) | |
| SE (Site Entry) | | LOSAP (LOS Access Point) | |
| PBAAE (PBA Access Entry, animals only) | | CRP (Carcasses Removal Pathways) | |
| DCD (Designated Cleaning and Disinfection vehicle station) | | VM (Vehicle Movements) | |
| GCD (Generalized Cleaning and Disinfection temporary station) | | LC (Loading Chute) | |
| PCD (Proposed Cleaning and Disinfection-temporary station) | | PBA (Perimeter Buffer Area) | |
| ADU (Dumpster for dead animals) | | LOS (Line of Separation) | |
| TD (Trash Dumpster) | | DPA (Designated Parking Area) | |
| SA (Supply drop-off Area) | | CD (Carcasses Disposal location/ADU box) | |

**Figure S1. Fabricated farm premises map containing farm features, including the Line of Separation (LOS) in red.** This represents an example of the maps developed for farms included in the Secure Pork Supply plans.

(A)

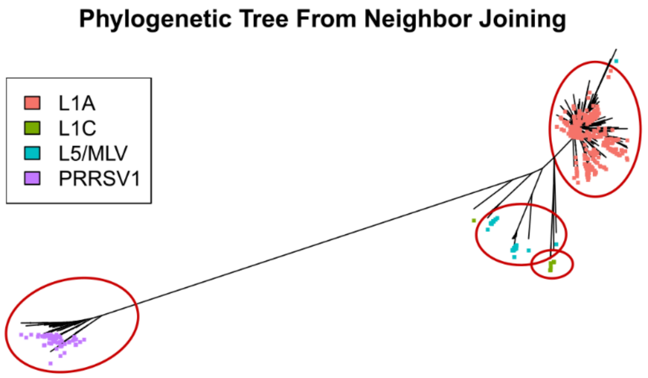


(B)

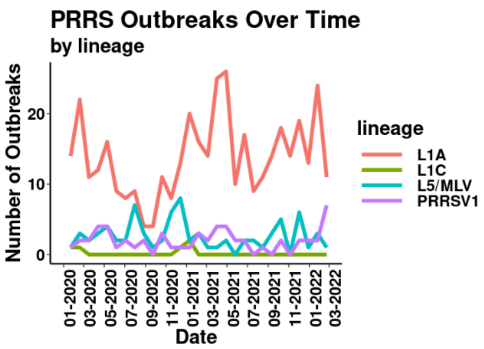


**Supplementary Figure S2:** Results of phylogenetic analysis of PRRSV data from January 2020 to March 2022. (A) shows the phylogenetic tree constructed with $R^2 = 0.98$. (B) shows the number of cases of each lineage identified.

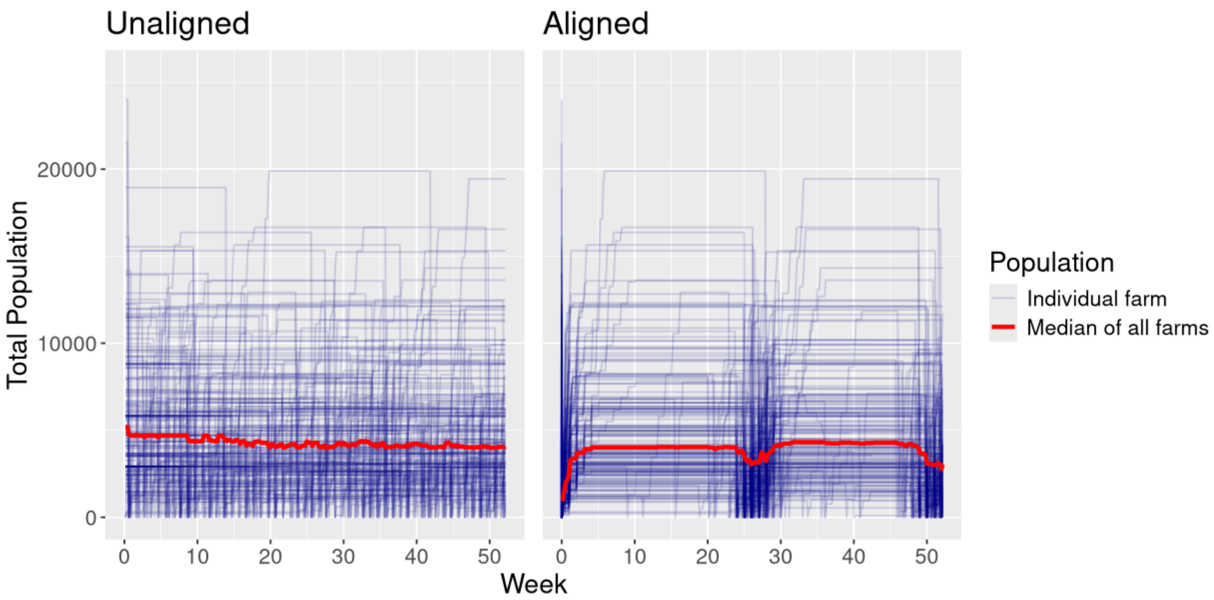


**Supplementary Figure S3:** The panels show the total population over time for each farm along with the median value of all farms. The curves are either unaligned or aligned as indicated with reference to the first restocking event during the period of July 1, 2022-December 31, 2023.

(A)

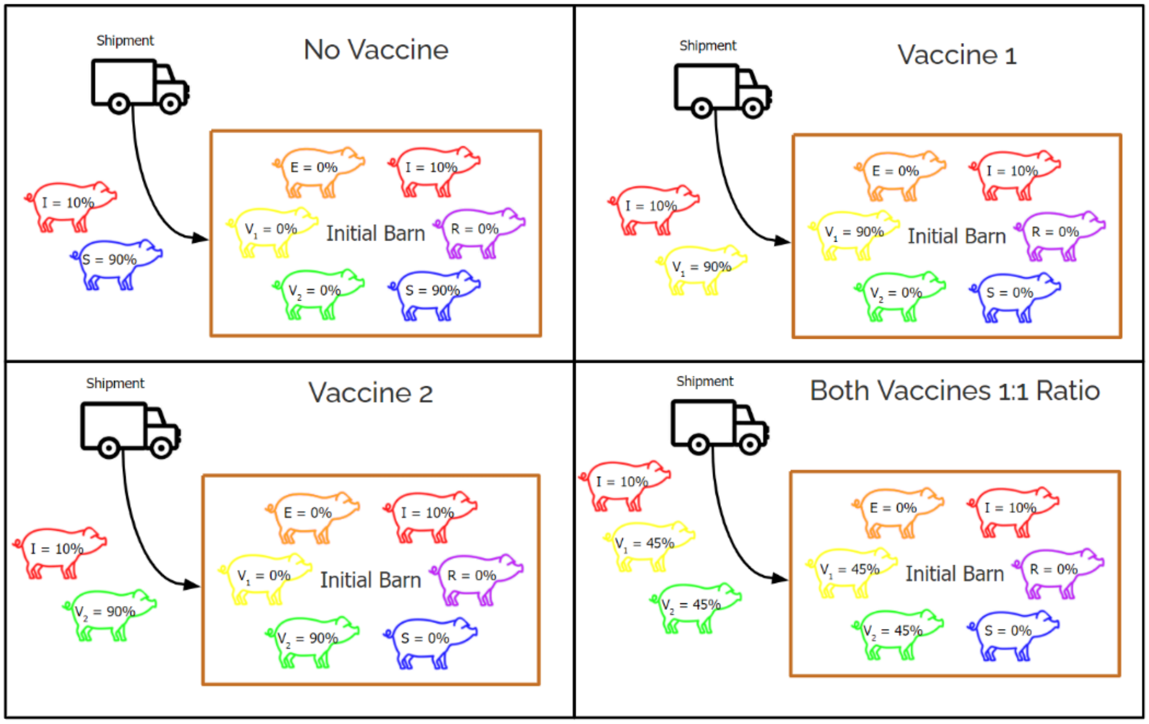
Shipment
No Vaccine
I = 10%
S = 90%
E = 0%
I = 10%
$V_1$ = 0%
Initial Barn
R = 0%
$V_2$ = 0%
S = 90%
Shipment
Vaccine 1
I = 10%
$V_1$ = 90%
E = 0%
I = 10%
$V_1$ = 90%
Initial Barn
R = 0%
$V_2$ = 0%
S = 0%
Shipment
Vaccine 2
I = 10%
$V_2$ = 90%
E = 0%
I = 10%
$V_1$ = 0%
Initial Barn
R = 0%
$V_2$ = 90%
S = 0%
Shipment
Both Vaccines 1:1 Ratio
I = 10%
$V_1$ = 45%
$V_2$ = 45%
E = 0%
I = 10%
$V_1$ = 45%
Initial Barn
R = 0%
$V_2$ = 45%
S = 0%

(B)

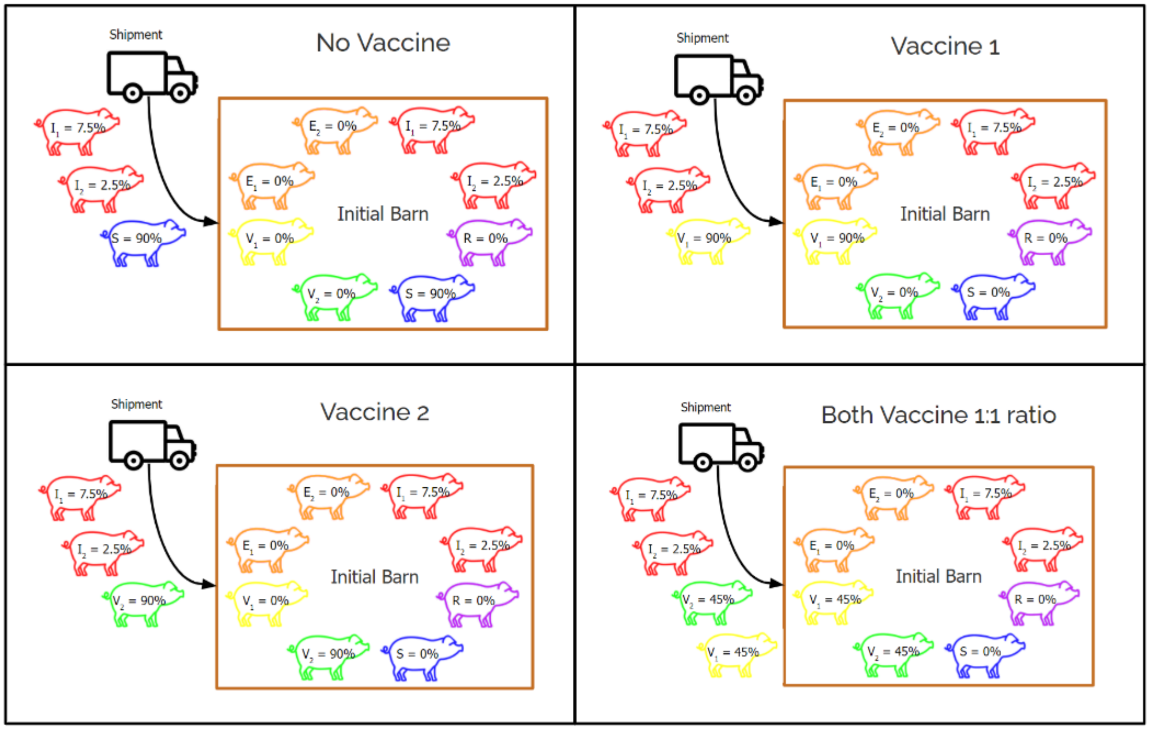
Shipment
No Vaccine
$I_1$ = 7.5%
$I_2$ = 2.5%
S = 90%
$E_2$ = 0%
$I_1$ = 7.5%
$E_1$ = 0%
$I_2$ = 2.5%
Initial Barn
$V_1$ = 0%
R = 0%
$V_2$ = 0%
S = 90%
Shipment
Vaccine 1
$I_1$ = 7.5%
$I_2$ = 2.5%
$V_1$ = 90%
$E_2$ = 0%
$I_1$ = 7.5%
$E_1$ = 0%
$I_2$ = 2.5%
Initial Barn
$V_1$ = 90%
R = 0%
$V_2$ = 0%
S = 0%
Shipment
Vaccine 2
$I_1$ = 7.5%
$I_2$ = 2.5%
$V_2$ = 90%
$E_2$ = 0%
$I_1$ = 7.5%
$E_1$ = 0%
$I_2$ = 2.5%
Initial Barn
$V_1$ = 0%
R = 0%
$V_2$ = 90%
S = 0%
Shipment
Both Vaccine 1:1 ratio
$I_1$ = 7.5%
$I_2$ = 2.5%
$V_2$ = 45%
$V_1$ = 45%
$E_2$ = 0%
$I_1$ = 7.5%
$E_1$ = 0%
$I_2$ = 2.5%
Initial Barn
$V_1$ = 45%
R = 0%
$V_2$ = 45%
S = 0%

**Supplementary Figure 4:** Diagrams illustrate the initial state of the barns and compartmental makeup of each incoming shipment. (A) shows the setup for the 1 strain scenarios and (B) shows the setup for the 2 strain scenarios.

(A)

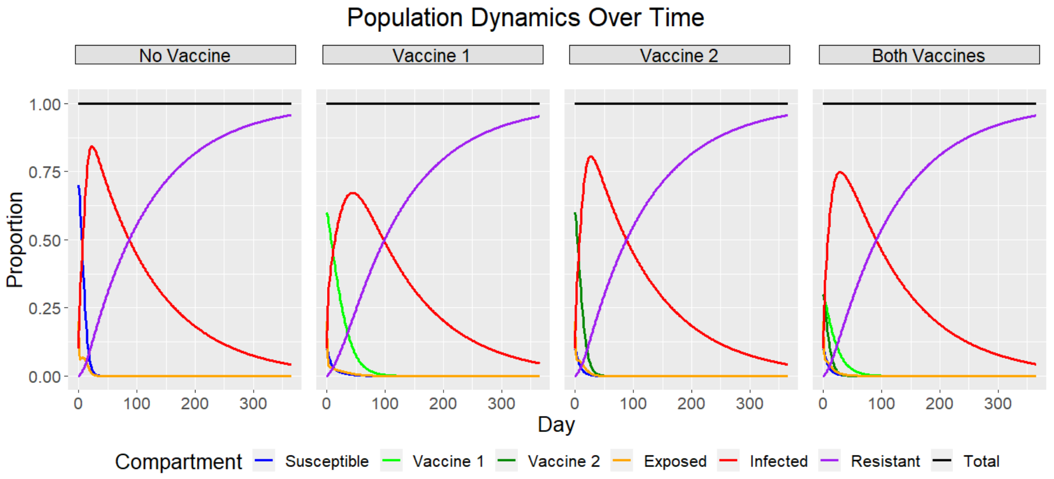


(B)

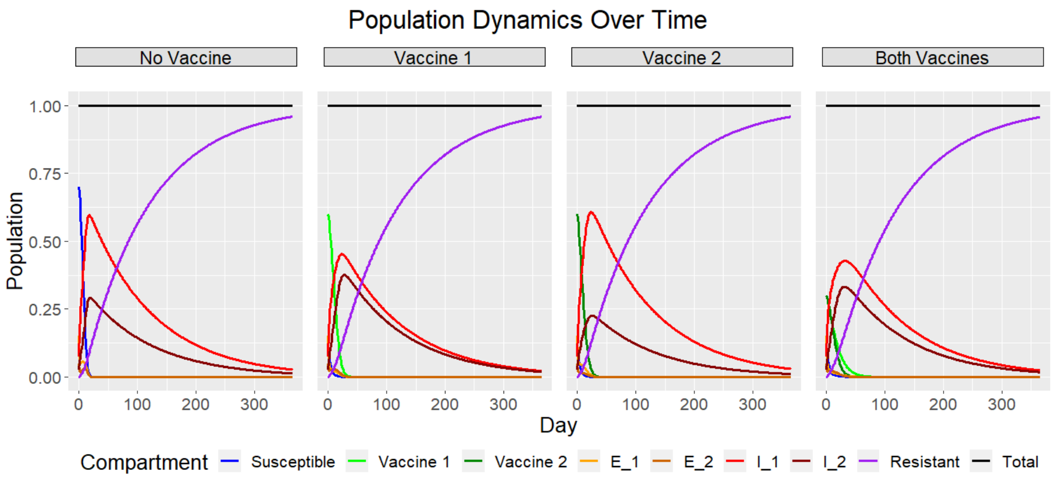


**Supplementary Figure 5:** Model dynamics for a fixed population of 1,000 over the course of a year. A) shows the dynamics for 1 strain, B) shows the 2 strain dynamics
